# Wheat photosystem II heat tolerance responds dynamically to short and long-term warming

**DOI:** 10.1101/2021.11.01.466822

**Authors:** Bradley C. Posch, Julia Hammer, Owen K. Atkin, Helen Bramley, Yong-Ling Ruan, Richard Trethowan, Onoriode Coast

## Abstract

Heat-induced inhibition of photosynthesis is a key factor in declining wheat performance and yield. Variation in wheat heat tolerance can be characterised using the critical temperature (*T*_crit_) above which incipient damage to the photosynthetic machinery occurs. We investigated intraspecies variation and plasticity of wheat *T*_crit_ under elevated temperature in field and controlled environment experiments. We also assessed whether intraspecies variation in wheat *T*_crit_ mirrors patterns of global interspecies variation in heat tolerance reported for mostly wild, woody plants. In the field, wheat *T*_crit_ varied through the course of a day, peaking at noon and lowest at sunrise, and increased as plants developed from heading to anthesis and grain filling. Under controlled temperature conditions, heat stress (36°C) was associated with a rapid rise in wheat *T*_crit_ (i.e. within two hours of heat stress) that peaked after 3–4 days. These peaks in *T*_crit_ indicate a physiological limitation to photosystem II heat tolerance. Analysis of a global dataset (comprising 183 *Triticum* and wild wheat (*Aegilops*) species) generated from the current study and a systematic literature review showed that wheat leaf *T*_crit_ varied by up to 20°C (about two-thirds of reported global plant interspecies variation). However, unlike global patterns of interspecies *T*_crit_ variation which has been linked to latitude of genotype origin, intraspecific variation in wheat *T*_crit_ was unrelated to that. Yet, the observed genotypic variation and plasticity of wheat *T*_crit_ suggests that this trait could be a useful tool for high-throughput phenotyping of wheat photosynthetic heat tolerance.

## Introduction

As the climate changes, global mean land-surface temperature has continued to rise, alongside more frequent, longer, and more intense heatwaves (Perkins-Kirkpatrick and Lewis, 2020). This is particularly concerning for the prospect of improving crop yields, as heat stress is associated with significant declines in the yield of widely-cultivated crops, including wheat (Asseng *et al*., 2015; Tack *et al*., 2015; Hochman *et al*., 2017; Liu *et al*., 2019; Ortiz-Bobea *et al*., 2019). Photosynthesis is a primary determinant of wheat yield and it is particularly sensitive to heat stress (Berry and Bjorkman, 1980; Way and Yamori, 2014). Improving the heat tolerance of photosynthesis could future-proof wheat yield in a warming world (Cossani and Reynolds, 2012; Scafaro and Atkin, 2016; Iqbal *et al*., 2017). To realise improvements in wheat photosynthetic heat tolerance, it is paramount that we first understand and quantify patterns of wheat photosynthetic heat tolerance so that we can then successfully exploit them.

Decreased leaf photosynthetic rate under high temperature is partially linked to disruption of the chloroplast electron transport chain, of which the thylakoid membrane-embedded photosystem II (PSII) considered the most sensitive component (Sharkey, 2005; Brestic *et al*., 2012). Heat-induced reactive oxygen species and lipid peroxidation both cause cleavage of the reaction centre-binding D1 protein of PSII (Yamashita *et al*., 2008), inhibiting electron flow and thus the production of ATP. For decades PSII damage has been measured with chlorophyll *a* fluorescence metrics, including *T*_crit_ of F_0_ (Schreiber *et al*., 1975; Schreiber and Berry, 1977; Hüve *et al*., 2011; Geange *et al*., 2021). *T*_crit_ of F_0_ (henceforth *T*_crit_) is the critical temperature above which minimal chlorophyll *a* fluorescence (F_0_) rises rapidly, indicating incipient damage to PSII (Schreiber and Berry, 1977; Melcarek and Brown, 1979; Neuner and Pramsohler, 2006; Slot *et al*., 2019). *T*_crit_ is associated with increased thylakoid membrane fluidity, disruption of the light-harvesting antennae (Raison *et al*., 1982; Figueroa *et al*., 2003), dissociation of chloroplast membrane-bound proteins (Berry and Bjorkman, 1980), and loss of chloroplast thermostability (Armond *et al*., 1978). As a standardized metric, *T*_crit_ has been used to examine global patterns of heat tolerance, quantify phenotypic plasticity in response to warming, and assess vulnerability to climate change across plant species (O’Sullivan *et al*., 2017; Zhu *et al*., 2018; Lancaster and Humphreys, 2020; Geange *et al*., 2021). While the number of publications examining plant *T*_crit_ is growing (Ferguson *et al*., 2020; Arnold *et al*., 2021; Slot *et al*., 2021), most studies focused on woody, non-crop species, and characterisation of intraspecies variation in *T*_crit_ of crop species has been limited (see Ferguson *et al*., 2020for a recent exception). Wheat, as the most widely-cultivated crop (with over 220 million ha cultivated worldwide) with a diverse range of genotypes originating from across the globe, is an ideal crop species for examining intraspecies variation and acclimation of *T*_crit_. In addition, although wheat is a temperate crop, there is increasing evidence of warming in many wheat-producing regions, including China, the USA and Australia, resulting in either stalled or reduced wheat yield (Hochman *et al*., 2017; Zhao *et al*., 2017). Understanding the response of *T*_crit_ to warming and the magnitude of intraspecies variation in *T*_crit_ could thus provide opportunities for improving photosynthetic heat tolerance and yield resilience in wheat and other crops.

Quantification of intraspecific variation in physiological traits of crops commonly encounters bottlenecks at the phenotyping stage. However, high-throughput phenotyping techniques are being developed, including a robotic system offering a ten-fold increase in the measurement speed of dark respiration (Scafaro *et al*., 2017; Coast *et al*., 2019, 2021), and the proximal remote sensing of leaf hyperspectral reflectance signatures for rapidly assaying photosynthetic characteristics and dark respiration (Silva-Pérez *et al*., 2018; Coast *et al*., 2019; Fu *et al*., 2019). Though chlorophyll fluorescence techniques are well-established for assessing photosynthetic heat tolerance, they are typically cumbersome. This limits their incorporation in breeding programmes that screen hundreds of genotypes for heat tolerance. But recently, Arnold *et al*. (2021) described a high-throughput chlorophyll fluorescence screening technique for a diversity of wild species. Previous studies of photosynthetic thermal tolerance also largely assumed that *T*_crit_ is diurnally and phenologically constant. However, these assumptions may be flawed. Substantial changes in metabolic capacity and demand for photosynthetic products vary diurnally, with phenological development and in response to fluctuations in temperature (Steer, 1973; Rashid *et al*., 2020). Thus, it seems reasonable that *T*_crit_ may demonstrate similar variation. However, these assumptions remain untested.

The extent to which plants physiologically adjust to warming is important in determining productivity and survival (Scheiner, 1993; Leung *et al*., 2020). Acclimation of photosynthetic electron transport to elevated temperature is evidenced by an increase in *T*_crit_. Zhu et al. (2018) reported acclimation at a rate of 0.34° C increase in *T*_crit_ for every 1°C increase in average temperature over the growing season for a range of native Australian species. Acclimation of *T*_crit_ may also increase plant thermal safety margins, thus protecting against damage to PSII under future heat stress. Thermal safety margins are estimated as the difference between the upper limit of leaf function (e.g. *T*_crit_) and the maximum growth temperature experienced in an environment (Sastry and Barua, 2017), and they provide a useful representation of a species’ potential vulnerability to global warming (Sunday *et al*., 2014). A reduction in this margin indicates increasing vulnerability to heat stress, while an increase in this margin indicates better capacity to tolerate the effects of climate warming (Hoffmann *et al*., 2013). Thermal safety margins of 10–15°C have been reported for many plant species (Weng and Lai, 2005; O’Sullivan *et al*., 2017; Perez and Feeley, 2020), with some as high as 12–31°C (Leon-Garcia and Lasso, 2019) when leaf temperature, rather than air temperature, was used. However, many plant species have low thermal safety margins (e.g. ≤ 5°C; Sastry and Barua, 2017). Unfortunately, reports quantifying the acclimation capacity and thermal safety margins of food crops are scarce. Reports on acclimation of *T*_crit_ to warming have been in response to a sustained increase in long-term growth temperature. Similar descriptions of *T*_crit_ acclimation to short-term heat stress (e.g. heatwaves) are not well documented. Considering heatwaves are predicted to become more frequent and intense (Perkins-Kirkpatrick and Lewis, 2020), it is pertinent that we understand if and how *T*_crit_ responds to heatwaves. Whether acclimation of *T*_crit_ to heatwaves has an upper threshold (i.e. a ceiling temperature) is currently unknown.

Previous uses of *T*_crit_ to assess global patterns of heat tolerance have been underpinned by ecological theories established in terrestrial ectotherms and endotherms (Addo-Bediako *et al*., 2000; Deutsch *et al*., 2008; Sunday *et al*., 2011; Araújo *et al*., 2013). One such theory is that organism physiology correlates closely with large-scale geographical patterns in the thermal environment where populations of an individual species were evolved (Gabriel and Lynch, 1992). Indeed, greater photosynthetic heat tolerance of non-crop plants originating from hotter, equatorial environments has been reported for numerous species (O’Sullivan *et al*., 2017; Drake *et al*., 2018; Zhu *et al*., 2018; Lancaster and Humphreys, 2020). It remains unknown whether such global patterns of interspecies variation hold for intraspecific comparisons – e.g. in a widely-cultivated crop like wheat, with genotypes originating from across the globe.

In this study, we employed a high-throughput system to describe intraspecies variation and high temperature acclimation of *T*_crit_ in wheat. Our objectives were to: (i) examine whether leaf *T*_crit_ varies diurnally and with phenology; (ii) determine the thermal safety margins and assess vulnerability of wheat to high temperatures in the Australian grain belt; and (iii) to assess if there is an upper threshold for leaf *T*_crit_ exposed to a sustained heat shock. To achieve these objectives, we conducted three field studies and one controlled environment experiments. In addition, we conducted a systematic literature review of wheat *T*_crit_ and used the global data we generated to investigate if intraspecies variation in wheat leaf *T*_crit_ is related to the latitude of genotype (as a proxy for climate of origin) of wheat genotypes or species.

## Materials and Methods

### Field experiments: Assessing diel and phenological variation in wheat T_crit_ and estimating thermal safety margins of Australian wheat

### Germplasm

A set of 20–24 wheat genotypes (Table S1) were used in three field experiments conducted in Australia across three years. Twenty of these genotypes were used in Coast *et al*. (2021) to assess acclimation of wheat photosynthesis and respiration to warming in two of the fields. The genotypes included: commercial Australian cultivars; heat tolerant materials developed by the centres of the Consultative Group on International Agricultural Research (CGIAR) in Mexico and Morocco and tested in warm areas globally; materials derived from targeted crosses between adapted hexaploid cultivars and heat tolerant Mexican hexaploid landraces, tetraploid emmer wheat (*T. dicoccon* Schrank ex Schübl.), Indian cultivars and synthetic wheat derived by crossing *Aegilops tauchii* with modern tetraploid durum wheat. All genotypes evaluated were hexaploid and chosen for their contrasting heat tolerance under high temperature conditions in Sudan (Gezira; 14.9°N 33°E), Australia (Narrabri, NSW; 30.27°S, 149.81°E) and Mexico (Ciudad Obregón; 27.5°N, 109.90°W).

### Experimental design and husbandry

The first two years of field experiments were undertaken in regional Victoria (2017, Dingwall; and 2018, Barraport West), and the third was in regional New South Wales (2019, Narrabri). A detailed description of the experimental designs for the 2017 and 2018 experiments are reported in Coast *et al*. (2021). Briefly, a diverse panel of genotypes were sown on three dates each in 2017 (20 genotypes) and 2018 (24 genotypes) to expose crops to different growth temperatures at a common developmental stage. The first time of sowing (TOS) for both experiments were within the locally recommended periods for sowing (early May). Subsequent sowing times were one month apart in June and July. Experiments were sown in three adjacent strips, one for each TOS. Each strip consisted of four replicate blocks. The 2019 field experiment was similar in all aspects to the 2018 field experiment, except for the following: (i) only two times of sowing were incorporated in the design; and (ii) the sowing times were approximately two months apart (17 May 2019 and 15 July 2019). Of the 24 genotypes sown in 2018 only 20, which were common to the 2017 and 2019 experiments, were assessed for *T*_crit_. All three field experiments were managed following standard agronomic practices for the region by the Birchip Cropping Group (www.bcg.org.au) in regional Victoria, and staff of the IA Watson Grains Research Centre at The University of Sydney and AGT, in Narrabri. A summary of the field experiments is presented in Table 1.

**Table 1.** Information on field experiments in Australia

| Experiment location and year | Year | Genotypes studied<sup>a</sup> | Mean daily maximum temperature at anthesis (°C) | Radiation (μmol photons m<sup>-2</sup> s<sup>-1</sup>)<sup>b</sup> |
| --- | --- | --- | --- | --- |
| Dingwall, Victoria | 2017 | 20 | 21.4–31.6 | 1394–1934 |
| Barraport West, Victoria | 2018 | 20 | 22.8–33.4 | 1706–2331 |
| Narrabri, New South Wales | 2019 | 24 | 22.6–34.1 | 1823–1950 |
| Experiment objective | TOS<sup>c</sup> | Genotypes studied | Brief description of method |  |
| *Diurnal variation in $T_{crit}$* |  |  |  |  |
| Dingwall, Victoria | 3 | 6 | Flag leaf $T_{crit}$ determined at anthesis at four consecutive time points occurring every six hours over an 18-hour period (i.e. solar noon, sunset, midnight, and sunrise). | |
| *Phenological variation in $T_{crit}$* |  |  |  |  |
| Barraport West, Victoria | 1–3 | 4 | Flag leaf $T_{crit}$ determined at heading, anthesis, and grain filling on the same day at 10 am from all three time of sowing plots. | |
| *Rate of acclimation of $T_{crit}$<sup>+d</sup> and calculation of thermal safety margins* |  |  |  |  |
| Dingwall, Victoria | 1–3 | 20 | Times of sowing varied so that plants sown later experienced warmer growth environments at a common developmental stage. Flag leaf $T_{crit}$ determined at anthesis. Thermal safety margins were estimated as the difference between genotype mean flag leaf $T_{crit}$ at anthesis and the maximum recorded air temperature at Dingwall/Barraport West (40°C) or Narrabri (40.8°C) in October (typical month of peak wheat anthesis). | |
| Barraport West, Victoria | 1–3 | 20 |  |  |
| Narrabri, New South Wales | 1–2 | 24 |  |  |

<sup>a</sup>Twenty genotypes were common to all experiments. The designation of all genotypes used in this study is provided in Supplementary Table S1;
<sup>b</sup>Mean maximum photosynthetically active radiation measured with Licor 6400XTs light sensors; <sup>c</sup>Time of sowing, where the first time of sowing was within the locally recommended sowing window, with subsequent times of sowing separated in one month intervals at Victoria, or two months at New South Wales; <sup>+d</sup>an additional $T_{crit}$ high temperature acclimation study was conducted under controlled environments with two of the 24 genotypes.

### Diel measurements of wheat T_crit_

Six of the 20 genotypes in the 2017 field experiment at Dingwall, Victoria were used to investigate diel variation. The six genotypes were two commercial cultivars (Mace and Suntop) and four breeding lines (with reference numbers 143, 2254, 2267, and 2316). These were chosen because they are representative of the diversity of the set of 20 genotypes (Coast *et al*., 2021). To determine if *T*_crit_ varied diurnally, flag leaves were harvested at anthesis (Zadok GS60-69; Zadoks *et al*., 1974) from plants of TOS3 at four consecutive time points occurring every six hours over an 18–hour period (solar noon, sunset, midnight, and sunrise).

### Phenological measurements of wheat T_crit_

A subset of four genotypes from the 20 in the 2018 field experiment at Barraport West, Victoria was used to assess phenological variation in *T*_crit_. The four genotypes were the breeding lines 2062, 2150, 2254, and 2267, the latter two of which were also used for diel measurements as described in the previous paragraph. Plants at heading (Zadok GS50–59), anthesis (Zadok GS60–69), and grain filling (Zadok GS70–79) were respectively chosen from fields of the three times of sowing. Flag leaves were harvested from the tallest tillers of plants at the different phenological stages at 10 am on the same day and used to determine *T*_crit_.

### Estimation of thermal safety margin of Australian wheat

All 20 genotypes in the 2017 and 2018 field experiments in Dingwall and Barraport West respectively, as well as all 24 genotypes in the 2019 field experiment in Narrabri were used to estimate thermal safety margins. Thermal safety margins were estimated as the difference between individual genotype *T*_crit_ and the maximum recorded air temperature at either Dingwall or Narrabri in October. Similar definitions of thermal safety margins as the difference between the measured temperature at which a species experiences irreversible physiological damage and the maximum measured temperature of the species’ habitat have been used in studies of animal ectotherms and plants (Deutsch *et al*., 2008; Sunday *et al*., 2014; O’Sullivan *et al*., 2017; Sastry and Barua, 2017). We considered Barraport West and Dingwall together for their historical weather records as they are in close proximity to one another in the Mallee district of Victoria, Australia. Weather data for Dingwall and Barraport West were obtained from the Australian Bureau of Meteorology covering the period 1910–2020. We used 40°C and 40.8°C (the maximum recorded air temperature for October) in Dingwall/Barraport West and Narrabri, respectively, to quantify thermal safety margins under current climatic scenarios. October was chosen as the upper threshold of exposure of field plants at anthesis to heat because all the later sown plants in this study were at anthesis in October. For future climatic conditions we added 2.6°C and 5°C to the current maximum temperature, with the 2.6°C addition representing the top end of the intermediate emission Representative Concentration Pathway (RCP) 4.5 IPCC scenario predicted for Eastern Australia by 2090 (1.3–2.6°C), and the 5°C addition similarly representing the top end of the high emission RCP 8.5 IPCC scenario predicted for Eastern Australia by 2090 (2.8–5°C; Climate Change Australia, 2021). Across all times of sowings in the three field experiments, flag leaves were harvested at anthesis (Zadok GS60-69) at a standardised time of 9–10 am to determine *T*_crit_ and estimate thermal safety margins.

### Controlled environment experiment: speed of acclimation and upper limit of leaf T_crit_ during heat shock

A controlled environment experiment was conducted to determine the speed and threshold of the response of *T*_crit_ to a sudden heat shock. Two wheat genotypes – 29 and 2267 (Table S1) – which contrasted in *T*_crit_ under common conditions were used to assess the speed and potential threshold of the response of wheat leaf *T*_crit_ to sudden heat shock. This experiment was conducted at the Controlled Environment Facilities of the Australian National University (ANU), Canberra, Australia.

### Plant husbandry

Seeds were germinated on saturated paper towel in covered plastic containers under darkness for one week. Germinated seedlings were planted in 1.05 L pots (130 mm diameter) filled with potting mix (80% composted bark, 10% sharp sand, 10% coir) with 4g L^-1^ fertiliser (Osmocote Exact Mini fertiliser, ICL, Tel Aviv, Israel) mixed through.

### Temperature treatment

Potted plants were grown in glasshouses in which a 24/18° C day/night temperature regime with a 12-hour photo-thermal period was maintained until tillering. At tillering, when all plants had a fully-extended third leaf (Zadok growth scale 22– 29; Zadoks *et al*., 1974), plants were moved into growth cabinets (TPG-2400-TH, Thermoline Scientific, Wetherill Park, NSW, Australia) for temperature treatment. One of two temperature conditions were imposed: a day/night regime of 24/18° C, or a heat shock with day/night temperatures of 36/24° C. White fluorescent tubes provided a 12 h photoperiod of photosynthetically active radiation of 720–750 µmol m^-2^ s^-1^ at plant height. Leaf discs were sampled from fully extended third leaves from main tillers and used to determine *T*_crit_ after 2, 4, 24, 48, 72, and 120 hours in the growth cabinets. Four replicate plants were used for *T*_crit_ measurement at each sampling time and for each temperature condition. Plant husbandry followed standard practice at the ANU Controlled Environment Facilities.

### Meta-study (field experiments, glasshouse studies and a systematic literature review) of wheat T_crit_ relationship with origins of genotypes

To explore how our results, compare with previous studies that have assessed wheat leaf *T*_crit_, and whether genotypes from hot habitats exhibit higher *T*_crit_, we undertook a systematic review of the published literature and compiled data from over 30 years (1988 to 2020) of wheat leaf *T*_crit_ studies. A database was generated using information from a recently published systematic review on global plant thermal tolerance (Geange *et al*., 2021) and additional literature search. These published data were combined with data from the three field experiments described above. The multiple times of sowings in each of the three Australian field experiments provided us with eight thermal environments for obtaining wheat leaf *T*_crit_ from a total of 24 wheat genotypes. We also included unpublished wheat leaf *T*_crit_ data from nine other experiments conducted in controlled environment facilities at the Australian National University. Overall, our global dataset included 3223 leaf *T*_crit_ samples from 183 wheat genotypes of various species (*T. aestivum L*., *T. turgidum* L., ssp. durum Desf., *T turgidum* L., ssp. diococcoides Thell.) and wild wheat (*Aegilops* species).

### Determination of leaf T_crit_

Leaves were harvested and stored in plastic bags alongside a saturated paper towel and were left to dark adapt for a minimum of 20 minutes. Water (90 μl) was placed in each well of a 48-well Peltier heating block in order to ensure leaf samples remained hydrated throughout the assay. A single 6 mm diameter leaf disc was excised from the middle of each harvested dark-adapted leaf and placed within each well of the heating block. Once discs were all loaded into the heating block, a glass plate was used to enclose the wells to prevent leaf pieces from drying out during the assay. The block was then placed directly beneath the lens of an imaging fluorometer (*FluorCam 800MF*, Photon Systems Instruments, Brno, Czech Republic) and programmed to heat from 20 to 65° C at a rate of 1°C min^-1^. The fluorometer recorded F_0_ throughout the heating period (approximately one record per minute). Following the conclusion of the temperature ramp, fluorescence data was processed and used to estimate *T*_crit_, which was calculated by fitting linear regressions to both the flat and the steep parts of the F_0_ curves and then recording the temperature at which these two lines intersected.

### Statistical analysis

Statistical analyses were carried out within the R statistical environment (v. 3.4.4; R Core Team, 2018) with R Studio. For analysis of the field data on diurnal and phenological variation in *T*_crit_ we employed linear mixed models in R using the packages lmerTest (Kuznetsova *et al*., 2017) and emmeans (Lenth, 2020). In the models we treated genotype and time of day (for the diel variation) or developmental stage (for the phenological variation) as fixed terms and replicate as a random term. The ceiling threshold of *T*_crit_ under heat stress (of 36°C), in the controlled environment experiment, was determined by fitting a non-linear regression to the *T*_crit_ by time relationship. Then using the coefficients of the fitted regressions, we estimated the time at which the fitted *T*_crit_ was highest and this was taken as the time to peak acclimation. To test the relationships between *T*_crit_ and growth environment temperature, we only used data from the three field experiments in Australia, for which we had reliable data. The 24 genotypes studied under field conditions in Australia were grouped based on the region of origin of their pedigree (Aleppo, Syria; Gezira, Sudan; Narrabri, Australia; Obregón, Mexico; Pune, India; and Roseworthy, Australia) and the relationship examined using linear or bivariate regressions. Our global dataset (see Table 2 for sources) was used to ascertain the link between wheat leaf *T*_crit_ and climate of origin by regressing mean genotype *T*_crit_ with genotype latitude (as a proxy for climate) of origin.

**Table 2.** Source of data used for assessment of global variation in leaf photosynthetic heat tolerance (T_crit_).

| Study<sup>1</sup> | Origin | Species | Mean $T_{crit}$ (n) |
| --- | --- | --- | --- |
| This study | Asia | *Triticum aestivum* L. | 45.1 (8) |
|  | Africa | *T. aestivum* L. | 44.6 (1) |
|  | Australia | *T. aestivum* L. and *T. dicoccum* Schrank | 44.7 (9) |
|  | North America | *T. aestivum* L. | 45.0 (6) |
| **Average** |  |  | **44.8 (24)** |
| Havaux *et al.* (1988) | Africa | *T. turgidum* L., ssp. *durum* Desf. | 49.7 (9) |
|  | Europe | *T. turgidum* L., ssp. *durum* Desf. | 48.1 (19) |
|  | North America | *T. turgidum* L., ssp. *durum* Desf. | 51.8 (1) |
|  | South America | *T. turgidum* L., ssp. *durum* Desf. | 49.0 (2) |
| **Average** |  |  | **48.7 (31)** |
| Végh *et al.* (2018) | Europe | *T. aestivum* L. | **41.8 (5)** |
| Rekika *et al.* (1997) | Africa | *T. turgidum* L., ssp. *durum* Desf. | 37.0 (1) |
|  | North America | *T. turgidum* L., ssp. *durum* Desf. | 35.0 (1) |
|  | Europe | *T. turgidum* L., ssp. *durum* Desf. | 37.2 (3) |
|  |  | *T. turgidum* L., ssp. *diococcoides* Thell. | 38.0 (1) |
|  | Europe (wild wheat) | *Aegilops species* | 38.2 (5) |
| **Average** |  |  | **37.5 (11)** |
| Unpublished data from our group | Asia | *T. aestivum* L. | 43.8 (21) |
|  | Africa | *T. aestivum* L. | 45.2 (1) |
|  | Australia | *T. aestivum* L. and *T. dicoccum* Schrank | 44.5 (79) |
|  | North America |  | 43.9 (32) |
| **Average** |  |  | **44.2 (133)** |

<sup>1</sup>The fluorescence temperature response curves used in these studies were similar (ramp rate of 1–1.5°C min<sup>-1</sup>, in the 20–65°C range). Values in bold are study averages and those in parentheses indicate number of genotypes/species used.

## Results

### Diel and phenological variation in T_crit_

There was significant genotype by time of day interaction for *T*_crit_ (*P*=0.042; Table S2), highlighting the heterogeneity in this diel variation of *T*_crit_ among our genotypes. In all but genotype 2316, *T*_crit_ tended to be highest at solar noon before then declining through sunset, midnight, and sunrise. The slope of these trends was only significant for genotype 2267, with *T*_crit_ declining by 3.1°C from solar noon to sunrise. By contrast, genotypes 143 exhibited the narrowest diel range in *T*_crit_, with difference of 1.1°C between solar noon and sunrise. Irrespective of genotype, *T*_crit_ at solar noon was significantly higher than at sunrise (*P*<0.001 for time of day). *T*_crit_ also showed significant genotype by phenology interaction, and highly significant differences for the main effects of genotype as well as phenology (Table S2). The interaction effect was largely due to the increasing trend in *T*_crit_ as plants developed from heading to anthesis and grain filling for genotypes 2267, 2254 and 2062 but not for 2150 (Fig. 1B). *T*_crit_ of genotype 2150 rose slightly between heading and anthesis then declined significantly at grain filling relative to anthesis. Genotype 2254 showed the largest increase in *T*_crit_ between heading and anthesis, rising 1.8°C from 44.4°C to 46.2°C.

**Figure 1.**
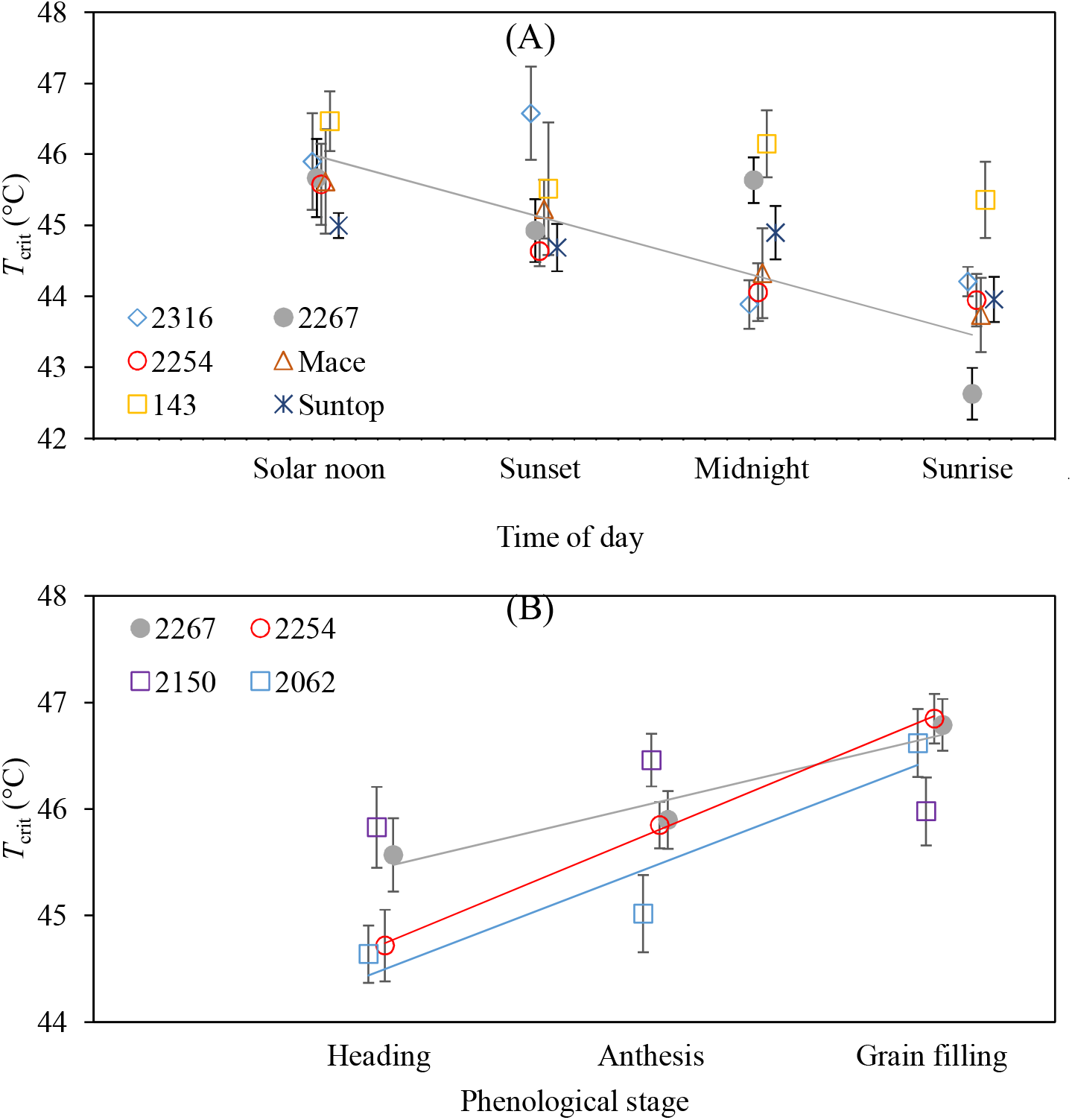
Variation in flag leaf *T*_crit_ (° C) of wheat genotypes over the course of an 18 h period (A), and across three phenological stages (B). Solid lines indicate significant linear trends. Plants were grown at field sites in Dingwall, Victoria in 2017 (A), and in Barraport West, Victoria in 2018 (B). Points represent mean ± se, *n* = 4 for (A) and *n* = 8–18 for (B).

### Thermal safety margins of Australian wheat

At Dingwall, only the main effect of TOS was significant (Table S3). In comparison to TOS 1, *T*_crit_ of TOS 2 tended to be higher and TOS 3 tended to be lower (Fig. 2A). At Barraport, *T*_crit_ varied amongst the genotypes over a range of about 2°C and the effect of TOS on *T*_crit_ depended on the genotype (*P*<0.01 for Genotype by TOS interaction). *T*_crit_ increased more in some genotypes (e.g. 1132, 143 and Trojan) under later sowing than others (e.g. 2267 and 29), but also did not change significantly in some (e.g. 1898 and 1943; Fig. 2B). At Narrabri, only the main effects were significant, i.e. genotypes varied in their *T*_crit_ and TOS 3 *T*_crit_ was higher than TOS 1 (Table S3, Fig. 2C). Across TOS, the genotypes with the lowest and highest mean *T*_crit_ were 1704 (45.3°C) and 143 (46.7°C) respectively. Across the 24 genotypes, *T*_crit_ increased by 0.5°C from TOS 1 (at 45.7°C) to TOS 3 (at 46.2°C). An analysis of variance run on a linear mixed effects model of the entire field data set revealed field site to be the largest source of variation in *T*_crit_ of all our independent variables (d.f. = 2, *F* value = 190.9, *P*<0.001). The overall mean *T*_crit_ at each of the field sites was 45.1°C at Barraport West, 44.1°C at Dingwall, and 45.9°C at Narrabri.

**Figure 2.**
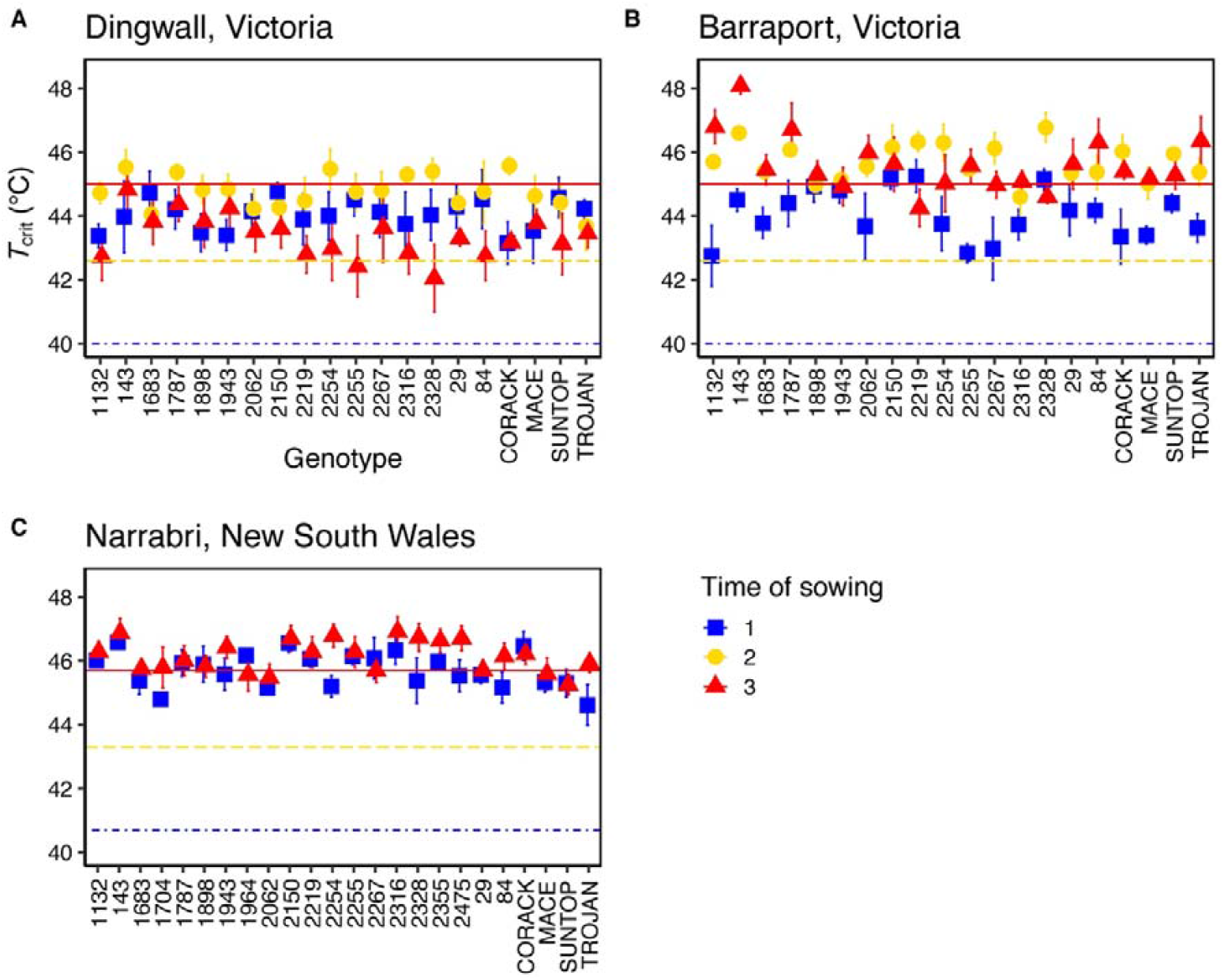
Phenotypic plasticity of leaf *T*_crit_ and thermal safety margins of 20–24 wheat genotypes. The genotypes were sown at either the locally recommended time of year (time of sowing 1; blue squares); one month after the recommended time (time of sowing 2; yellow circles); or two months after the recommended time (time of sowing 3; red triangles) at three Australian field sites. Delayed times of sowing were used to impose warmer average growth temperatures for plants sown at times of sowing 2 and 3. The field sites were Dingwall (A) and Barraport West (B), Victoria, and Narrabri, New South Wales (C). Twenty genotypes were sown at Dingwall in 2017 and Barraport West in 2018, and the same 20 plus an additional four genotypes were sown at Narrabri in 2019. The dash-dot blue lines mark the hottest recorded maximum temperature during the typical anthesis month (October) at each field site (40.7°C for Narrabri, and 40°C for Dingwall, data from the Australian Bureau of Meteorology; due to the close proximity of Dingwall and Barraport West we used the same climate records for these sites) while the yellow dashed line and the red solid line mark the RCP 4.5 IPCC and RCP 8.5 IPCC emission scenarios (+2.6 and +5°C), respectively. The difference between the observed *T*_crit_ and these current and future maximum temperatures is termed the thermal safety margin. Here we assume that leaf temperature is equal to air temperature. Points represent mean ± s.e., minimum *n* = 4.

Thermal safety margins were calculated for all field-grown genotypes by quantifying the difference between *T*_crit_ and the maximum air temperature recorded during October. All genotypes demonstrated a higher *T*_crit_ than the historical maximum October air temperatures recorded at each field site (Fig. 2, dash-dot blue line). Thermal safety margins in the TOS 1 fields ranged from 3.2–4.8°C in Dingwall (Fig. 2A), to 2.8–5.3°C in Barraport West (Fig. 2B) and from 3.8–5.8°C in Narrabri (Fig. 2C). For the later grown crops (i.e. TOS 2 and 3) which experienced warmer growth temperatures, thermal safety margins increased relative to TOS 1 in Dingwall (3.7–5.6°C for TOS 2), in Barraport West (4.6–6.8°C for TOS 2, and 4.3–8.1°C for TOS 3) and in Narrabri (4.5–6.1°C). The exception to this pattern was TOS 3 at Dingwall where the lower end of the thermal safety margin range declined, resulting in a range of 2.1– 4.8°C. At both Narrabri and Barraport West, mean *T*_crit_ of all genotypes was above the +2.6°C mark associated with the RCP 4.5 intermediate emission scenario (Fig. 2B & 2C, dashed yellow line). Most genotypes were also largely clear of the RCP 4.5 mark at Dingwall, except the *T*_crit_ of genotypes 2255 and 2328 sown at TOS 3 were below this threshold. The +5°C warming mark associated with the high emission RCP 8.5 scenario was equal to or above the *T*_crit_ of many genotypes at all three field sites, though there was some variation across the locations. At Narrabri, half of the genotypes were below the RCP 8.5 threshold when sown at TOS 1, while this fell to a quarter of genotypes when sown at TOS 3 (Fig. 2C). At TOS 1 in Barraport West, 17 genotypes fell below the RCP 8.5 threshold, with only one and three genotypes below this mark for TOS 2 and 3, respectively (Fig. 2B). At Dingwall, *T*_crit_ of all genotypes sown at TOS 1 and TOS 3 was below the RCP 8.5 threshold, while 14 genotypes at TOS 2 were below this mark (Fig. 2A).

### Genotype origin does not predict variation or acclimation in T_crit_

The 24 genotypes grown across the three field sites were grouped by the regions from which they originated (Table S1; Aleppo, Syria; Gezira, Sudan; Narrabri, Australia; Obregón, Mexico; Pune, India; and Roseworthy, Australia) in order to determine if this explained any of the observed variation in *T*_crit_. Genotype origin had a significant effect on *T*_crit_ at both Barraport West and Narrabri (Table S4) At Barraport West, genotypes that originated in Narrabri had the highest *T*_crit_ and Sudan the lowest (Fig. 3B). At Narrabri, the genotype that originated from Syria had the highest *T*_crit_ whereas those from Roseworthy had the lowest. By contrast, genotype origin had no significant effect on *T*_crit_ at Dingwall. TOS had a significant effect on *T*_crit_ at all three sites irrespective of origin. *T*_crit_ was lower than TOS 3 relative to TOS 1/TOS 2 in Dingwall, and was higher than TOS 1 in TOS 2/TOS 3 in Barraport West (Fig. 3). At the Narrabri site, *T*_crit_ increased from TOS 1 to TOS 3 for all origin groupings (Fig. 3C). No interaction between time of sowing and genotype origin was observed at any field site (Table S4).

**Figure 3.**
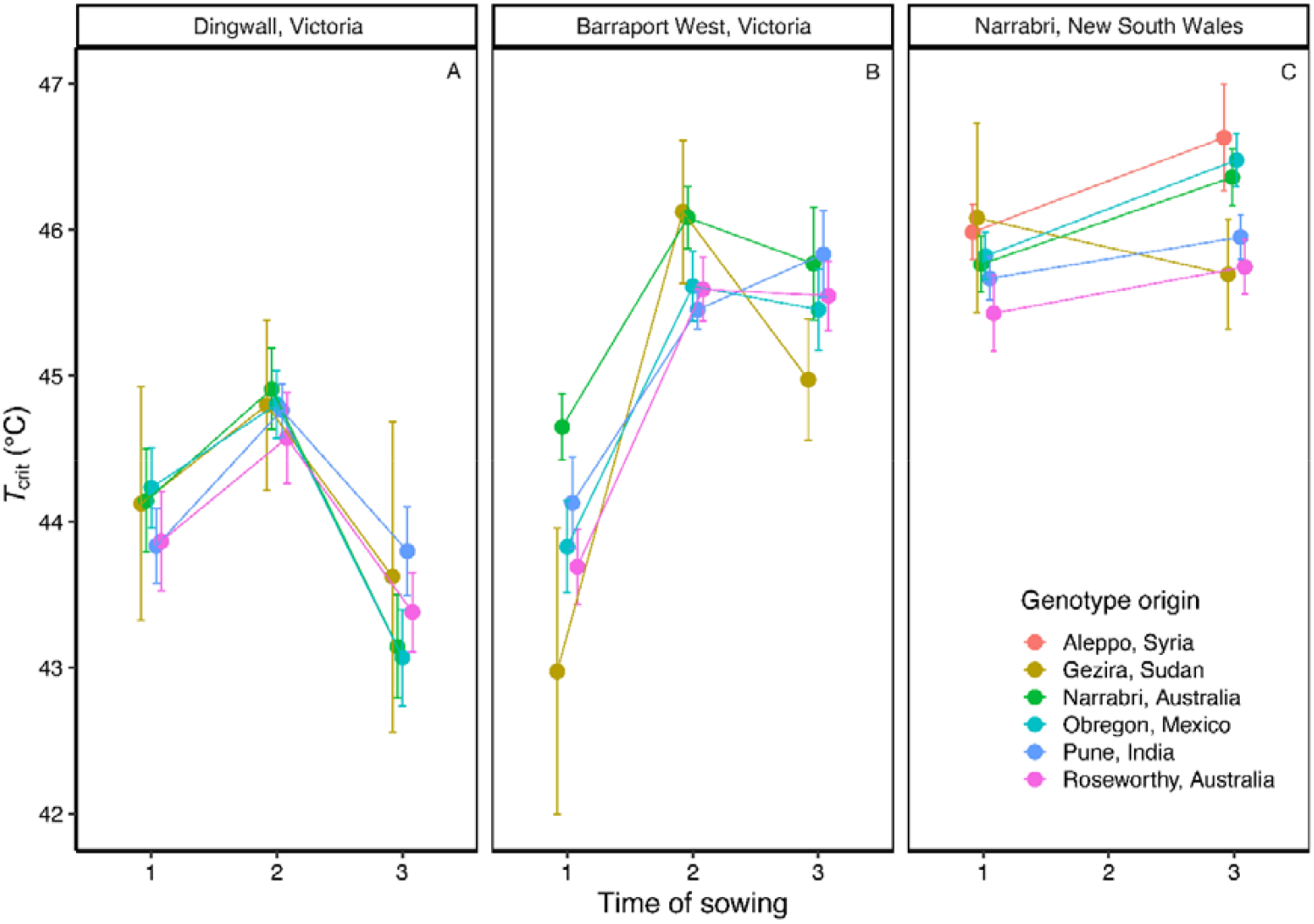
Phenotypic response of wheat flag leaf photosystem II heat tolerance (*T*_crit_) to time of sowing at three Australian field sites: (A) Dingwall, Victoria; (B) Barraport West, Victoria; and (C) Narrabri, New South Wales. Genotypes are grouped according to the six locations of the breeding programmes where they were developed. Twenty genotypes were grown at Dingwall in 2017 and at Barraport West in 2018, while the same 20 plus an additional four genotypes were grown in Narrabri in 2019. In order to generate increasingly warmer growth temperature regimes plants were sown at one of three times of sowing: time of sowing 1 (TOS 1) was the locally recommended time of sowing, while time of sowing 2 (TOS 2) and time of sowing 3 (TOS 3) were one and two months after TOS 1, respectively. Points represent mean ± se, minimum *n* = 4.

### Response of T_crit_ to short-term exposure to high temperature and upper limit of T_crit_ plasticity

In the two genotypes studied, *T*_crit_ increased significantly following two hours of heat shock (Fig. 4). In both genotypes, *T*_crit_ increased during the heat shock following a curvilinear pattern which peaked after 3.4 days for genotype 2267 and 4.2 days for genotype 54. Although the time to reach peak *T*_crit_ during the heat shock differed for the two genotypes, their maximum *T*_crit_ values were similar, being 43.8° C for genotype 29 and 43.6° C for genotype 2267 (Fig. 4). *T*_crit_ for both genotypes remained largely constant over the 120-hour period for those plants that were maintained at the control day/night temperature regime of 24/12°C.

**Figure 4.**
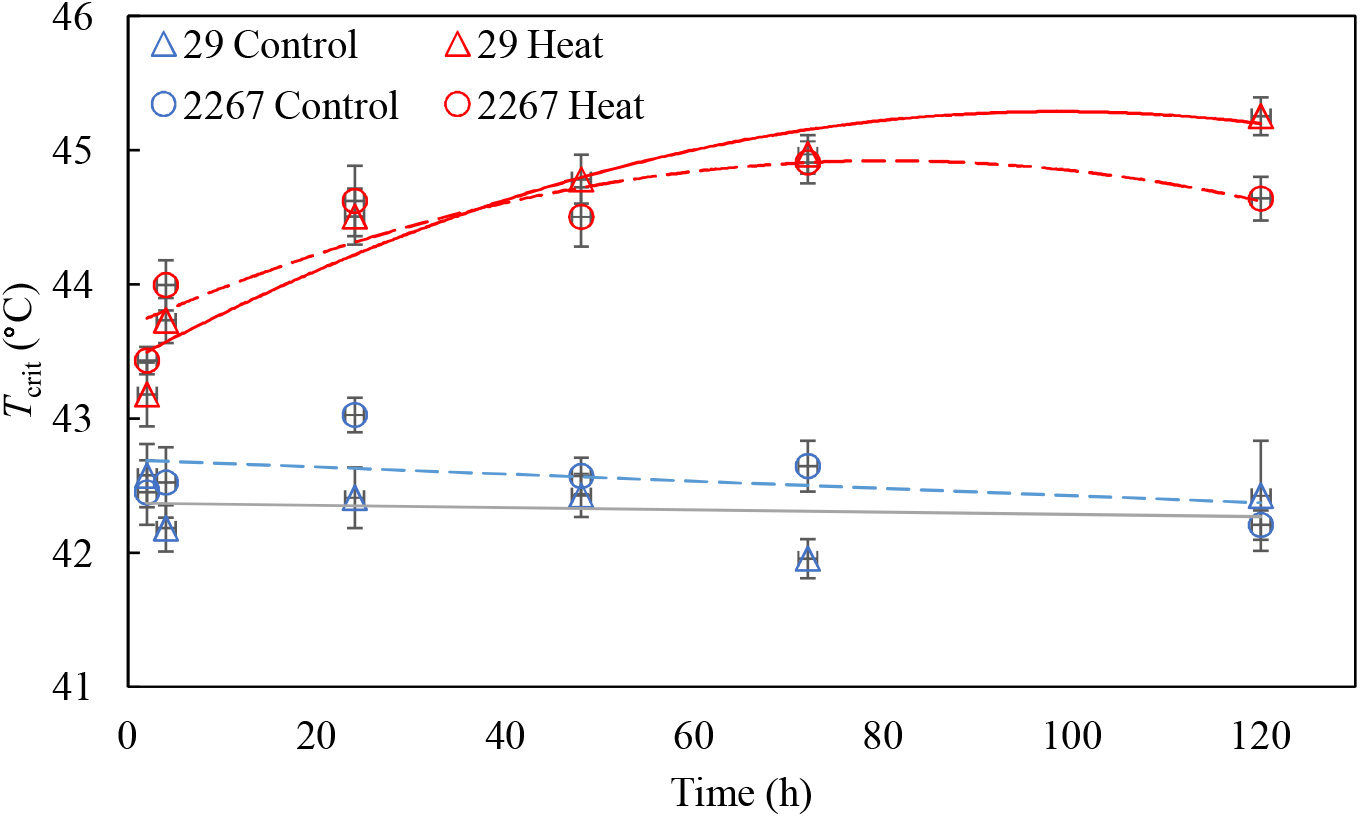
Leaf *T*_crit_ (° C) of two wheat genotypes – 29 (triangles and solid lines) and 2267 (circles and dashed lines) exposed to 24° C (control; blue shapes and lines) or 36° C (heat; red shapes and lines) for varying durations (2, 4, 24, 48, 72, or 120 h) in a growth cabinet. Leaf samples for *T*_crit_ were from the third fully extended leaves on the main stem. Equations for the curvilinear relationships between *T*_crit_ at 36° C (*T*_crit_ ^36^; ° C) and time (*t*; hour) under heat for genotype 29 is *T*_crit_ ^36^ = 43.42 + 0.038*t* – 0.00019*t*^2^ and for genotype 2267 is *T*_crit_ ^36^ = 43.69 + 0.031*t* – 0.00019*t*^2^. Points represent mean ± se, *n* = 4.

### Global variation in wheat T_crit_

We combined data from our experiments with previously published data (covering genotypes grown across field and controlled environment experiments) to examine the degree of variation in *T*_crit_ in wheat genotypes on a global scale based on the latitude of origin as a proxy for climate of origin of their pedigree (Fig. 5). We found three studies (Havaux *et al*., 1988; Rekika *et al*., 1997; Végh *et al*., 2018) that reported wheat leaf *T*_crit_ using similar fluorescence temperature response curves (with ramp rates of 1–1.5°C min^-1^ between 20– 65°C) to estimate *T*_crit_. Our final data collation comprised 183 wheat species/varieties (comprising *T. aestivum* L., *T. turgidum* L., ssp. *durum* Desf., *T. turgidum* L., ssp. *diococcoides* Thell., and wild wheat – *Aegilops* species) originating from all continents except Antarctica (Table 2). Globally, wheat leaf *T*_crit_ varied by up to 20°C (35–55°C) and there were more data for studies under warm conditions for genotypes originating from the lower latitudes than high latitudes (Fig 5). The larger variation in *T*_crit_ for genotypes originating from the higher latitudes coincided with the cooler growth conditions. Overall, there was less variation in *T*_crit_ under the warm conditions. We found no relationship between wheat leaf *T*_crit_ and the absolute latitude of genotype climate of origin (Fig. 5).

**Figure 5.**
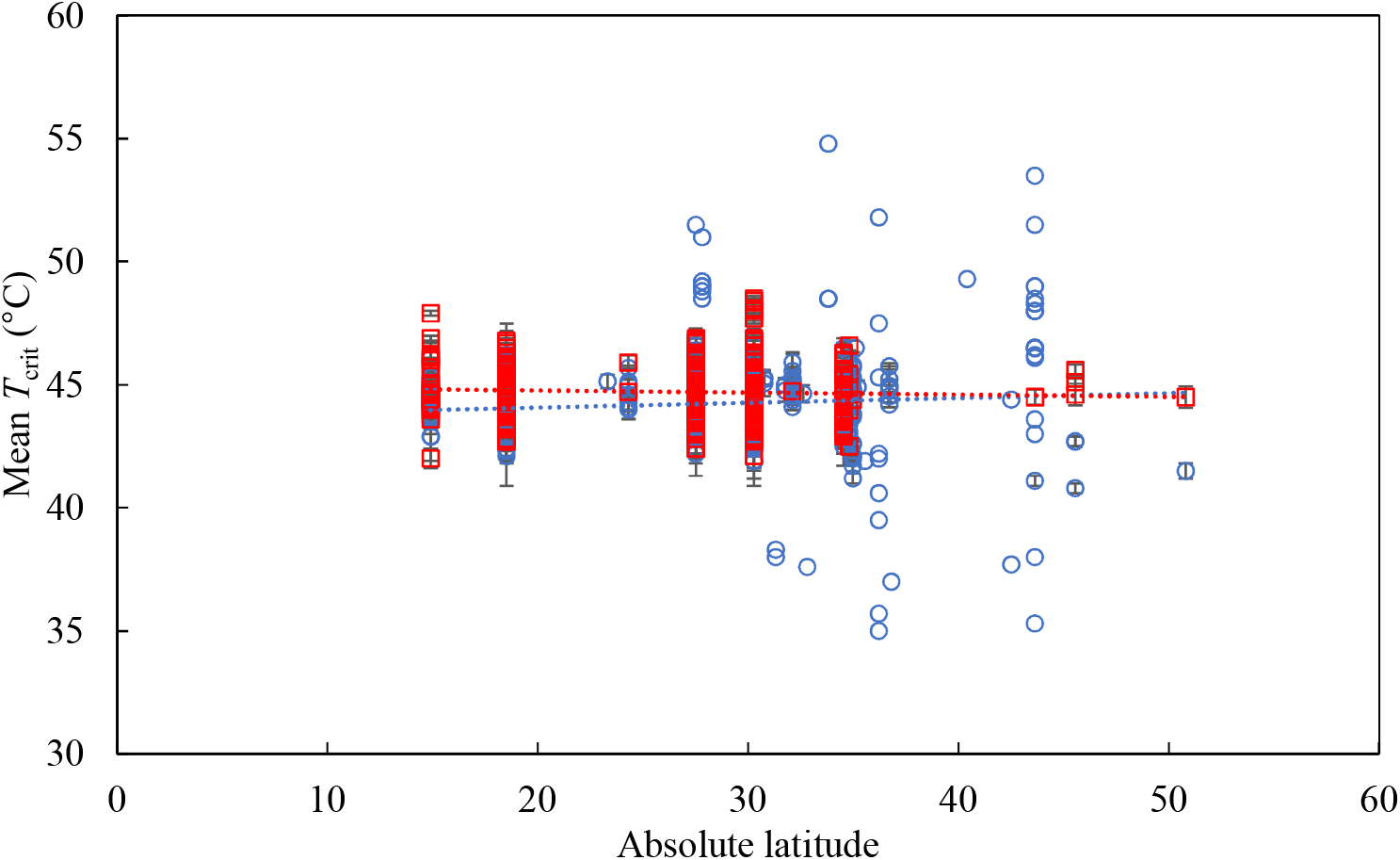
Relationship between *T*_crit_ and the absolute latitude of the climate of origin for wheat genotypes when grown under cool (blue circles) and warm (red squares) conditions.Data obtained from 183 wheat genotypes (3223 measurements of leaf *T*_crit_ overall) from experiments in Australia (this study) and published literature (Havaux *et al*., 1988; Rekika *et al*., 1997; Végh *et al*., 2018). Data points represent mean *T*_crit_ (± SE where visible) for each genotype.

## Discussion

In this paper we used a high-throughput technique to record minimal chlorophyll *a* fluorescence and quantified the critical temperature (*T*_crit_) of photosystem II damage – a measure of leaf photosynthetic heat tolerance – for wheat genotypes grown in multiple field experiments, as well as a controlled environment experiment. The field experiments demonstrated the extent of variation in *T*_crit_ over the course of a single day, as well as across several crucial stages of phenological development. They also showed that the region of origin of wheat genotypes were unrelated to *T*_crit_ in three representative Australian wheat growing regions, and that sowing time (and thus, growth temperature) was responsible for significant variation in *T*_crit_. Delayed sowing (i.e. elevated growth temperature) was generally associated with increases in *T*_crit_, resulting in higher thermal safety margins at both field sites. When two genotypes were subjected to a sudden heat shock in a controlled environment, we observed a slight difference between genotypes in the speed with which *T*_crit_ increased. However, both genotypes exhibited a similar peak *T*_crit_ value during this heat shock. Finally, when combining these data with previously published wheat *T*_crit_ data, as well as unpublished data from other experiments conducted in controlled environment facilities at The Australian National University, we found that the absolute latitude of pedigrees of wheat genotypes were not significantly linked with variation in *T*_crit_ for either cool or warm grown plants.

### Temporal fluctuations in wheat T_crit_ may be linked to changes in leaf sugar content

Wheat *T*_crit_ varied significantly over the course of a single day, declining by an average of 1.7° C over the 18 hours from solar noon to sunrise (Fig. 1A). This pattern resembles the extent of change in *T*_crit_ in a temperate tree species reported by Hüve *et al*. (2006); specifically, a linear increase over 14 hours, from a low point at 5 am to a peak at 7 pm. Taken together, these findings suggest that *T*_crit_ generally increases to a peak during the late afternoon before declining to a minimum between midnight and dawn. Hüve *et al*. (2006) linked this diel variation in *T*_crit_ with daily variation in leaf sugar content, and demonstrated that *T*_crit_ increased when leaves were fed sugar solutions. Further work is needed to determine if the diel variation in *T*_crit_ that we observed in wheat was also influenced by corresponding variation in leaf sugar content.

It is also interesting to compare the extent of variation in *T*_crit_ that was observed over the course of a single day with the extent of variation that was observed across phenology. In the 18 hours between solar noon and sunrise *T*_crit_ declined by 1.7°C (Fig. 1A), a fluctuation that was similar in size to the 1.5°C rise in *T*_crit_ that we observed from heading to anthesis (Fig. 1B). This comparison highlights the high level of plasticity in *T*_crit_, and that variation in *T*_crit_ is clearly responsive to factors on both an hours-long timescale (i.e. diurnal fluctuations in leaf sugar content) and a longer-term weeks-long scale (as evidenced by changes in *T*_crit_ from heading to anthesis and grain filling). Anthesis is widely considered the stage of phenology at which wheat is most vulnerable to high temperature (Ferris *et al*., 1998; Thistlethwaite *et al*., 2020), and so it was somewhat surprising to see *T*_crit_ rise as plants moved from heading to anthesis at the first glance. However, it is not surprising when considering the fact that anthesis vulnerability to heat stress is largely due to reduction of sink strength to import and utilize assimilates within the reproductive organs, rather than of assimilate supply from leaf photosynthesis per se (Li *et al*., 2012; Ruan *et al*., 2012). While this increase could reflect an ongoing rise in heat tolerance coinciding with seasonal warming, there was no significant difference in *T*_crit_ between plants undergoing anthesis versus those at the grain filling stage, and so anthesis may be the phenological stage at which *T*_crit_ is at its peak.

### Drivers of variation in wheat T_crit_

The field site at which plants were grown was the most significant source of variation in *T*_crit_; the overall average *T*_crit_ at Narrabri was 1.8°C higher than recorded at Dingwall and 0.8°C higher than at Barraport West. In addition to environment, genotype had significant effect on *T*_crit_ at the Barraport West and Narrabri sites. These results suggest that environment, genotype and most likely the genotype-by-environment interactions (GEI) may play large roles in determining wheat flag leaf *T*_crit_. Breeding for genotypes with greater photosynthetic heat tolerance (i.e. higher *T*_crit_) may be challenging if variation in *T*_crit_ is also influenced by GEI effects. GEI effects have been reported for other abiotic stress tolerance traits including lodging tolerance in spring wheat (Dreccer *et al*., 2020), and drought tolerance in maize (Dias *et al*., 2018).

### Genotypes maintain moderate photosynthetic thermal safety margins

We observed variation in the thermal safety margins of wheat genotypes, predominantly associated with differences between field sites and the effect of sowing time at these sites. The thermal safety margin was 2.1°C when averaged across all genotypes (Fig. 2). Thermal safety margins in three representative Australian wheat-growing regions were at least 2–4°C for all genotypes. *T*_crit_ was always several degrees greater than the hottest recorded air temperature during the typical month of anthesis at each site (Narrabri, 40.8°C, Dingwall/Barraport West 40°C; denoted by the blue dot-dash lines in Fig. 2). Under the IPCC’s RCP 4.5 intermediate emission scenario for Eastern Australia by 2090, most genotypes would maintain a positive, yet reduced, thermal safety margin in the studied growing regions. However, under the high emission RCP 8.5 scenario, the thermal safety margins of most genotypes grown at the Dingwall site and a few genotypes at the Barraport West site would be exceeded (Fig. 2A & 2B). For genotypes grown at the Narrabri site, thermal safety margins under the RCP 8.5 scenario would be drastically reduced and, in some cases, disappear (Fig. 2C). According to our *T*_crit_ observations, only genotypes originating from Obregón and Aleppo would retain positive thermal safety margins under the RCP 8.5 scenario when sown at either optimal or delayed sowing times. The rise in *T*_crit_ with delayed sowing (and thus increased growth temperature) that we observed in the majority of genotypes indicates a widespread capacity for the thermal acclimation of wheat flag leaf *T*_crit_. This suggests that thermal safety margins for wheat photosynthetic heat tolerance could yet increase in response to warming under future climate scenarios. However, given that we also observed an apparent limit to the acclimation of *T*_crit_ following sudden heat shock (Fig. 4), it is possible that daytime maximum temperatures could approach this physiological thermal limit of wheat PSII if the most severe global warming predictions are borne out. A hard limit to the high temperature acclimation of *T*_crit_ could indicate a physiological limitation of PSII, or a temperature that represents the absolute maximum tolerance. Given that the considerable thermal plasticity of PSII electron transport has been closely linked with improving photosynthetic heat tolerance more generally (Yamasaki *et al*., 2002), the prospect of air temperatures approaching the physiological threshold of PSII high temperature acclimation is concerning.

Thus far, in assessing thermal safety margins we have assumed parity between air and leaf temperatures; however, wheat leaf/canopy temperature can differ substantially from air temperature. Balota et al. (2007) reported canopy temperatures ranging from 3ºC below noon air temperatures to 10ºC above noon air temperatures in dryland wheat, and 3ºC below noon air temperatures to 5.7ºC above noon air temperatures in irrigated wheat. Similarly, canopy temperatures of Australian wheat have also been recorded exceeding afternoon air temperature by 0.3–2.3°C (Rattey *et al*., 2011) and 3–5°C (Rebetzke *et al*., 2013). These examples, along with other previous instances (Rashid *et al*., 1999; Thapa *et al*., 2018), highlight the significant genotypic variation in canopy cooling and thus the potential for achieving gains in performance under high temperature by exploiting this variation. While greater levels of canopy cooling could increase thermal safety margins by limiting leaf temperature, achieving gains in wheat *T*_crit_ could also provide an avenue to maintaining positive thermal safety margins by increasing the threshold to PSII damage. Enhancing thermal safety margins by increasing *T*_crit_ could be particularly important in water-limited environments considering that heatwaves are frequently accompanied by drought, which increases stomatal closure and limits transpirational cooling, resulting in increased leaf temperature (Aspinwall *et al*., 2019).

### Thermal environment of growth site may be more influential than genotype origin in determining variation in wheat flag leaf T_crit_

Considering the potential benefits to wheat heat tolerance and performance under high temperature that could arise from achieving increases in *T*_crit_, as well as the extent of variation that we observed in *T*_crit_ among 24 genotypes at three field sites, it would be beneficial to identify characteristics that predict high *T*_crit_ in wheat genotypes. Thus, we analysed whether the distinct regions from which our genotypes originated could reliably predict variation in *T*_crit_. Previous studies of (mostly) woody, non-crop species found that *T*_crit_ was correlated with climate of origin (O’Sullivan *et al*., 2017; Zhu *et al*., 2018). In a similar vein, we found evidence of genotype region of origin significantly affecting *T*_crit_ at two of our field sites (Fig. 3A & 3C). One consistency at both of these sites was that genotypes originating from Roseworthy, Australia generally exhibited the lowest or second-lowest mean *T*_crit_ values. By contrast, the genotype from Aleppo exhibited the highest *T*_crit_ at the Narrabri site (Fig. 3C), while at the Barraport West site the genotypes originating from Narrabri had the highest mean *T*_crit_ across all times of sowing (Fig. 3A). However, it seems unlikely that the effect of genotype region of origin is the result of differences in temperature at these locations: for instance, the average daily maximum April temperature in Aleppo, Syria is 23°C (NOAA), while the average daily October maximum in Roseworthy, Australia is 23.8°C (Australian Bureau of Meteorology). Therefore, the differences associated with genotype origin in the current study are likely related to a more complex combination of environmental differences between locations (e.g. rainfall, temperature, soil quality, agricultural practices). Differences in the aims and methods of breeding programs at various locations could also explain variation in *T*_crit_ associated with genotype origin. We also note that our experiments did not include genotypes that were developed in cooler environments, such as wheat growing regions in Europe or Northern America, and so further work may be required to capture the full extent of variation in *T*_crit_ that is associated with various wheat growing regions.

### T_crit_ increases within hours of heat shock, and peaks after 3–4 days

We observed widespread evidence of wheat *T*_crit_ plasticity following exposure to high temperature, including elevated growth temperature in the field (via delayed sowing, Fig. 2 & 3) and sudden heat shock under controlled conditions (Fig. 4). We also saw clear genotypic variation in the plasticity of *T*_crit_ across these experiments. In some genotypes *T*_crit_ rose by upwards of 4°C when sowing time was delayed by two months (Fig. 2B), while in others *T*_crit_ showed no change or even declined by up to 1.2°C from TOS 1 to TOS 3 (Fig. 2A). Similarly, following a heat shock imposed under controlled conditions we observed a difference between two genotypes in the speed at which *T*_crit_ increased despite the two genotypes eventually reaching a similar peak *T*_crit_ (Fig. 4). Genotypic variation is thus evident not only in wheat flag leaf *T*_crit_ under common non-stressful temperatures, but also in the extent of *T*_crit_ plasticity in response to sudden heat shock. Increases in *T*_crit_ with warming have been reported previously (O’Sullivan *et al*., 2017; Zhu *et al*., 2018) and these are considered examples of high temperature acclimation. That we observed similar patterns in wheat *T*_crit_, as well as genotypic variation in this acclimation, suggests that the capacity to increase PSII heat tolerance could be a trait worth targeting for the development of wheat genotypes with greater heat tolerance. However, further work is needed to first investigate whether such acclimation is associated with enhanced performance under high temperature in the field.

One aspect of the current study that may aid such future efforts is the development of high throughput minimal chlorophyll *a* fluorescence assays that can be used for large-scale screening of wheat PSII heat tolerance. When combined with other burgeoning high throughput techniques for measuring photosynthetic characteristics (Sharma *et al*., 2012; Silva-Pérez *et al*., 2018; Fu *et al*., 2019; McAusland *et al*., 2019; Arnold *et al*., 2021), it is becoming increasingly achievable to efficiently measure a range of traits that provide insight into the photosynthetic thermal tolerance of entire plots in crop breeding trials.

### Genotypic variation in wheat leaf T_crit_ is not consistent with latitudinal trends in general plant heat tolerance

Contrary to previous results that observed a decrease in PSII heat tolerance (measured as *T*_crit_) as latitude moved further from the equator (O’Sullivan *et al*., 2017; Lancaster and Humphreys, 2020), we found that, irrepective of thermal acclimation, wheat leaf *T*_crit_ did not vary with the latitude of genotype climate of origin (Fig. 5). This discrepancy could be related to differences between cultivated and wild species: the O’Sullivan *et al*. (2017) and Lancaster and Humphreys (2020) studies demonstrated a relationship between heat tolerance and latitude based almost entirely on records of different wild species. By contrast, our study focuses solely on one domesticated species. Wheat is known as a crop with a particularly narrow genetic background (Tanksley and McCouch, 1997), but we observed a large range of *T*_crit_ in wheat here (up to 20°C) which compares with the approximately 30°C global range reported across 218 plant species spanning seven biomes reported by O’Sullivan *et al*. (2017). This large range of wheat leaf *T*_crit_ can be exploited to improve heat tolerance in modern crop varieties, similar to recent successes in improving wheat drought tolerance using airborne remote sensing (Reynolds *et al*., 2015). Still, wheat is cultivated in a wide range of ecological and climatic conditions, covering over 220 million hectares, including areas where it is exposed to high temperature stress. As such, we predicted that the rise in *T*_crit_ that we observed with elevated growth temperature in our experimental data set (Fig. 1–4) would also be apparent in the meta-analysis. However, there was no evidence of any thermal acclimation response of *T*_crit_ in this larger data set. This could partly be due to diversity of experimental methods used to generate the data in Figure 5, as well as variation in the duration and intensity of elevated growth temperature treatments. Given that the plant thermal tolerance field uses a large and diverse range of experimental designs and assays (Geange *et al*., 2021), the results of our systematic review of wheat *T*_crit_ could be further evidence of a need to better standardise the approaches used to measure and describe photosynthetic heat tolerance.

## Conclusion

Wheat leaf *T*_crit_ varied dynamically with changes in growth conditions, notably increasing in response to short and long-term high temperatures, and exhibiting an upper ceiling in acclimating to heatwaves. There was also evidence of developmental, diel and genotypic variation in *T*_crit_. These results suggested a strong genotype-by-environment interaction effects on wheat leaf *T*_crit_ and potential links between *T*_crit_ and leaf sugar content. Interestingly, global wheat leaf *T*_crit_ which spanned up to 20°C was unrelated to genotype climate of origin and latitude, unlike reported associations with global interspecies variation in leaf *T*_crit_ of 171 plant species (*cf*. ∼30°C). However, the observed genotypic variation and plasticity of wheat *T*_crit_, combined with a recent developments of high throughput technique for measuring *T*_crit_ (Arnold et al 2021), indicate that this trait would be useful for high-throughput screening, understanding photosynthetic heat tolerance and development of heat tolerant wheat.

## Acknowledgements

We acknowledge and celebrate the First Australians on whose traditional lands this research was undertaken, and pay our respect to the elders past and present. This work was supported by grants from the ARC Centre of Excellence in Plant Energy Biology (CE140100008), and the Australian Grains Research and Development Corporation (GRDC) projects Postdoctoral Fellowship: Photosynthetic Acclimation to High Temperature in Wheat (US1904-003RTX – 9177346) and National Wheat Heat Tolerance (US00080) and Australian Research Council (DP180103834). Bradley C. Posch was supported by the Australian Government Research Training Program. Onoriode Coast also received support from Research England’s ‘Expanding Excellence in England’ (E3)-funded Food and Nutrition Security Initiative of the Natural Resources Institute, University of Greenwich. We are grateful to Claire Pickles and Amy Smith of Birchip Cropping Group, Victoria, for managing the trials in Victoria, and AGT and Sabina Yasmin for the trials in Narrabri. We are also grateful to the farmers who generously provided us with field sites for trials. Staff of the ANU Research School of Biology Plant Services team, especially Christine Larsen, Jenny Rath, Gavin Pritchard and Steven Dempsey are thanked for maintaining the plants in the controlled environments.

## References

Addo-Bediako A, Chown SL, Gaston KJ. 2000. Thermal tolerance, climatic variability and latitude. Proceedings of the Royal Society of London. Series B: Biological Sciences 267, 739–745.

Araújo MB, Ferri□Yáñez F, Bozinovic F, Marquet PA, Valladares F, Chown SL. 2013. Heat freezes niche evolution (D Sax, Ed.). Ecology Letters 16, 1206–1219.

Armond PA, Schreiber U, Björkman O. 1978. Photosynthetic Acclimation to Temperature in the Desert Shrub, Larrea divaricata. Plant Physiology 61, 411–415.

Arnold PA, Briceño VF, Gowland KM, Catling AA, Bravo LA, Nicotra AB. 2021. A high-throughput method for measuring critical thermal limits of leaves by chlorophyll imaging fluorescence. Functional Plant Biology 48, 634–646.

Aspinwall MJ, Pfautsch S, Tjoelker MG, et al.. 2019. Range size and growth temperature influence Eucalyptus species responses to an experimental heatwave. Global Change Biology 25, 1665–1684.

Asseng S, Ewert F, Martre P, et al.. 2015. Rising temperatures reduce global wheat production. Nature Climate Change 5, 143–147.

Balota M, Payne WA, Evett SR, Lazar MD. 2007. Canopy temperature depression sampling to assess grain yield and genotypic differentiation in winter wheat. Crop Science 47, 1518–1529.

Berry J, Bjorkman O. 1980. Photosynthetic response and adaptation to temperature in higher plants. Annual Review of Plant Physiology 31, 491–543.

Brestic M, Zivcak M, Kalaji HM, Carpentier R, Allakhverdiev SI. 2012. Photosystem II thermostability in situ: Environmentally induced acclimation and genotype-specific reactions in Triticum aestivum L. Plant Physiology and Biochemistry 57, 93–105.

Climate Change Australia. 2021. Climate change in Australia: Climate information, projections, tools and data. Available at: https://www.climatechangeinaustralia.gov.au/en/

Coast O, Posch BC, Bramley H, Gaju O, Richards RA, Lu M, Ruan Y, Trethowan R, Atkin OK. 2021. Acclimation of leaf photosynthesis and respiration to warming in field□grown wheat. Plant, Cell & Environment 44, 2331–2346.

Coast O, Shah S, Ivakov A, et al.. 2019. Predicting dark respiration rates of wheat leaves from hyperspectral reflectance. Plant, Cell & Environment 42, 2133–2150.

Cossani CM, Reynolds MP. 2012. Physiological traits for improving heat tolerance in wheat. Plant Physiology 160, 1710–1718.

Deutsch CA, Tewksbury JJ, Huey RB, Sheldon KS, Ghalambor CK, Haak DC, Martin PR. 2008. Impacts of climate warming on terrestrial ectotherms across latitude. Proceedings of the National Academy of Sciences of the United States of America 105, 6668–6672.

Dias KODG, Gezan SA, Guimarães CT, et al.. 2018. Estimating genotype × environment interaction for and genetic correlations among drought tolerance traits in maize via factor analytic multiplicative mixed models. Crop Science 58, 72–83.

Drake JE, Tjoelker MG, Vårhammar A, et al.. 2018. Trees tolerate an extreme heatwave via sustained transpirational cooling and increased leaf thermal tolerance. Global Change Biology 24, 2390–2402.

Dreccer MF, Condon AG, Macdonald B, et al.. 2020. Genotypic variation for lodging tolerance in spring wheat: wider and deeper root plates, a feature of low lodging, high yielding germplasm. Field Crops Research 258, 107942.

Feeley KJ, Martinez-Villa J, Perez T, Silva Duque A, Triviño Gonzalez D, Duque A. 2020. The thermal tolerances, distributions, and performances of tropical montane tree species. Frontiers in Forests and Global Change 3, 25.

Ferguson JN, McAusland L, Smith KE, Price AH, Wilson ZA, Murchie EH. 2020. Rapid temperature responses of photosystem II efficiency forecast genotypic variation in rice vegetative heat tolerance. Plant Journal 104, 839–855.

Ferris R, Ellis RHR, Wheeler TR, Hadley P. 1998. Effect of high temperature stress at anthesis on grain yield and biomass of field-grown crops of wheat. Annals of Botany 82, 631–639.

Figueroa FL, Conde-Álvarez R, Gomez I. 2003. Relations between electron transport rates determined by pulse amplitude modulated chlorophyll fluorescence and oxygen evolution in macroalgae under different light conditions. Photosynthesis Research 75, 259–275.

Fu P, Meacham-Hensold K, Guan K, Bernacchi CJ. 2019. Hyperspectral leaf reflectance as proxy for photosynthetic capacities: An ensemble approach based on multiple machine learning algorithms. Frontiers in Plant Science 10, 1–13.

Gabriel W, Lynch M. 1992. The selective advantage of reaction norms for environmental tolerance. Journal of Evolutionary Biology 5, 41–59.

Geange SR, Arnold PA, Catling AA, et al.. 2021. The thermal tolerance of photosynthetic tissues: a global systematic review and agenda for future research. New Phytologist 229, 2497–2513.

Havaux M, Ernez M, Lannoye R. 1988. Correlation between heat tolerance and drought tolerance in cereals demonstrated by rapid chlorophyll fluorescence tests. Journal of Plant Physiology 133, 555–560.

Hochman Z, Gobbett DL, Horan H. 2017. Climate trends account for stalled wheat yields in Australia since 1990. Global Change Biology 23, 2071–2081.

Hoffmann AA, Chown SL, Clusella-Trullas S. 2013. Upper thermal limits in terrestrial ectotherms: how constrained are theyã Functional Ecology 27, 934–949.

Hüve K, Bichele I, Rasulov B, Niinemets U. 2011. When it is too hot for photosynthesis: heat-induced instability of photosynthesis in relation to respiratory burst, cell permeability changes and H_2_O_2_ formation. Plant, Cell & Environment 34, 113–126.

Hüve K, Bichele I, Tobias M, Niinemets Ü. 2006. Heat sensitivity of photosynthetic electron transport varies during the day due to changes in sugars and osmotic potential. Plant, Cell & Environment 29, 212–228.

Iqbal M, Raja NI, Yasmeen F, Hussain M, Ejaz M, Shah MA. 2017. Impacts of heat stress on wheat: A critical review. Advances in Crop Science and Technology 5, 1–9.

Kuznetsova A, Brockhoff PB, Christensen RHB. 2017. lmerTest package: tests in linear mixed effects models. Journal of Statistical Software 82.

Lancaster LT, Humphreys AM. 2020. Global variation in the thermal tolerances of plants. Proceedings of the National Academy of Sciences of the United States of America 117, 13580–13587.

Lenth R. 2020. Estimated Marginal Means, aka Least-Squares Means. R package version 1.5.2-1.

Leon-Garcia I V., Lasso E. 2019. High heat tolerance in plants from the Andean highlands: Implications for paramos in a warmer world. PLOS ONE 14, e0224218.

Leung C, Rescan M, Grulois D, Chevin L. 2020. Reduced phenotypic plasticity evolves in less predictable environments. Ecology Letters 23, 1664–1672.

Li ZM, Palmer P, Martin M, Wang R, Rainsford F, Jin Y, Patrick JW, Yang YJ, Ruan Y-L. 2012 High invertase activity in tomato reproductive organs correlates with enhanced sucrose import into, and heat tolerance of, young fruit. Journal of Experimental Botany 63, 1155–1166.

Liu B, Martre P, Ewert F, et al.. 2019. Global wheat production with 1.5 and 2.0°C above pre-industrial warming. Global Change Biology 25, 1428–1444.

McAusland L, Atkinson JA, Lawson T, Murchie EH. 2019. High throughput procedure utilising chlorophyll fluorescence imaging to phenotype dynamic photosynthesis and photoprotection in leaves under controlled gaseous conditions. Plant Methods 15, 1–15.

Melcarek PK, Brown GN. 1979. Chlorophyll fluorescence monitoring of freezing point exotherms in leaves. Cryobiology 16, 69–73.

Neuner G, Pramsohler M. 2006. Freezing and high temperature thresholds of photosystem II compared to ice nucleation, frost and heat damage in evergreen subalpine plants. Physiologia Plantarum 126, 196–204.

O’Sullivan OS, Heskel MA, Reich PB, et al.. 2017. Thermal limits of leaf metabolism across biomes. Global Change Biology 23, 209–223.

Ortiz-Bobea A, Wang H, Carrillo CM, Ault TR. 2019. Unpacking the climatic drivers of US agricultural yields. Environmental Research Letters 14, 064003.

Perez TM, Feeley KJ. 2020. Photosynthetic heat tolerances and extreme leaf temperatures. Functional Ecology 34, 2236–2245.

Perkins-Kirkpatrick SE, Lewis SC. 2020. Increasing trends in regional heatwaves. Nature Communications 11, 1–8.

Raison JK, Roberts JKM, Berry JA. 1982. Correlations between the thermal stability of chloroplast (thylakoid) membranes and the composition and fluidity of their polar lipids upon acclimation of the higher plant, Nerium oleander, to growth temperature. BBA - Biomembranes 688, 218–228.

Rashid FAA, Scafaro AP, Asao S, Fenske R, Dewar RC, Masle J, Taylor NL, Atkin OK. 2020. Diel- and temperature-driven variation of leaf dark respiration rates and metabolite levels in rice. New Phytologist 228, 56–69.

Rashid A, Stark JC, Tanveer A, Mustafa T. 1999. Use of canopy temperature measurements as a screening tool for drought tool for drought tolerance in spring wheat. Journal of Agronomy and Crop Science 182, 231–238.

Rattey AR, Shorter R, Chapman SC. 2011. Evaluation of CIMMYT conventional and synthetic spring wheat germplasm in rainfed sub-tropical environments. II. Grain yield components and physiological traits. Field Crops Research 124, 195–204.

Rebetzke GJ, Rattey AR, Farquhar GD, Richards RA, Condon AG. 2013. Genomic regions for canopy temperature and their genetic association with stomatal conductance and grain yield in wheat. Functional Plant Biology 40, 14–33.

Rekika D, Monneveux E, Havaux M. 1997. The in vivo tolerance of photosynthetic membranes to high and low temperatures in cultivated and wild wheats of the Triticum and Aegilops genera. Journal of Plant Physiology 150, 734–738.

Reynolds M, Tattaris M, Cossani CM, Ellis M, Yamaguchi-Shinozaki K, Pierre C Saint. 2015. Exploring Genetic Resources to Increase Adaptation of Wheat to Climate Change. Advances in Wheat Genetics: From Genome to Field. Tokyo: Springer Japan, 355–368.

Ruan Y-L, Patrick JW, Bouzayen M, Osorio S, Fernie AR. 2012 Molecular regulation of seed and fruit set. Trends in Plant Science 17, 656–665.

Sastry A, Barua D. 2017. Leaf thermotolerance in tropical trees from a seasonally dry climate varies along the slow-fast resource acquisition spectrum. Scientific Reports 7, 11246.

Scafaro AP, Atkin OK. 2016. The Impact of Heat Stress on the Proteome of Crop Species. Agricultural Proteomics Volume 2. Cham: Springer International Publishing, 155–175.

Scafaro AP, Negrini ACA, O’Leary BM, et al.. 2017. The combination of gas-phase fluorophore technology and automation to enable high-throughput analysis of plant respiration. Plant Methods 13, 1–13.

Scheiner SM. 1993. Genetics and evolution of phenotypic plasticity. Annual Review of Ecology and Systematics 24, 35–68.

Schreiber U, Berry JA. 1977. Heat-induced changes of chlorophyll fluorescence in intact leaves correlated with damage of the photosynthetic apparatus. Planta 136, 233–238.

Schreiber U, Colbow K, Vidaver W. 1975. Temperature–jump chlorophyll fluorescence induction in plants. Zeitschrift für Naturforschung C 30, 689–690.

Sharkey TD. 2005. Effects of moderate heat stress on photosynthesis: Importance of thylakoid reactions, rubisco deactivation, reactive oxygen species, and thermotolerance provided by isoprene. Plant, Cell & Environment 28, 269–277.

Sharma DK, Andersen SB, Ottosen CO, Rosenqvist E. 2012. Phenotyping of wheat cultivars for heat tolerance using chlorophyll a fluorescence. Functional Plant Biology 39, 936–947.

Silva-Pérez V, Molero G, Serbin SP, Condon AG, Reynolds MP, Furbank RT, Evans JR. 2018. Hyperspectral reflectance as a tool to measure biochemical and physiological traits in wheat. Journal of Experimental Botany 69, 483–496.

Slot M, Cala D, Aranda J, Virgo A, Michaletz ST, Winter K. 2021. Leaf heat tolerance of 147 tropical forest species varies with elevation and leaf functional traits, but not with phylogeny. Plant, Cell & Environment, 2414–2427.

Slot M, Krause GH, Krause B, Hernández GG, Winter K. 2019. Photosynthetic heat tolerance of shade and sun leaves of three tropical tree species. Photosynthesis Research 141, 119–130.

Steer BT. 1973. Diurnal variations in photosynthetic products and nitrogen metabolism in expanding leaves. Plant Physiology 51, 744–748.

Sunday JM, Bates AE, Dulvy NK. 2011. Global analysis of thermal tolerance and latitude in ectotherms. Proceedings of the Royal Society B: Biological Sciences 278, 1823–1830.

Sunday JM, Bates AE, Kearney MR, Colwell RK, Dulvy NK, Longino JT, Huey RB. 2014. Thermal-safety margins and the necessity of thermoregulatory behavior across latitude and elevation. Proceedings of the National Academy of Sciences 111, 5610–5615.

Tack J, Barkley A, Nalley LL. 2015. Effect of warming temperatures on US wheat yields. Proceedings of the National Academy of Sciences of the United States of America 112, 6931–6936.

Tanksley SD, McCouch SR. 1997. Seed banks and molecular maps: Unlocking genetic potential from the wild. Science 277, 1063–1066.

Thapa S, Jessup KE, Pradhan GP, Rudd JC, Liu S, Mahan JR, Devkota RN, Baker JA, Xue Q. 2018. Canopy temperature depression at grain filling correlates to winter wheat yield in the U.S. Southern High Plains. Field Crops Research 217, 11–19.

Thistlethwaite RJ, Tan DKY, Bokshi AI, Ullah S, Trethowan RM. 2020. A phenotyping strategy for evaluating the high-temperature tolerance of wheat. Field Crops Research 255, 107905.

Végh B, Marček T, Karsai I, Janda T, Darkó É. 2018. Heat acclimation of photosynthesis in wheat genotypes of different origin. South African Journal of Botany 117, 184–192.

Way DA, Yamori W. 2014. Thermal acclimation of photosynthesis: On the importance of adjusting our definitions and accounting for thermal acclimation of respiration. Photosynthesis Research 119, 89–100.

Weng JH, Lai MF. 2005. Estimating heat tolerance among plant species by two chlorophyll fluorescence parameters. Photosynthetica 43, 439–444.

Yamasaki T, Yamakawa T, Yamane Y, Koike H, Satoh K, Katoh S. 2002. Temperature acclimation of photosynthesis and related changes in photosystem II electron transport in winter wheat. Plant Physiology 128, 1087–1097.

Yamashita A, Nijo N, Pospíšil P, Morita N, Takenaka D, Aminaka R, Yamamoto Y, Yamamoto Y. 2008. Quality control of photosystem II: Reactive oxygen species are responsible for the damage to photosystem II under moderate heat stress. Journal of Biological Chemistry 283, 28380–28391.

Zhao C, Liu B, Piao S, et al.. 2017. Temperature increase reduces global yields of major crops in four independent estimates. Proceedings of the National Academy of Sciences of the United States of America 114, 9326–9331.

Zhu L, Bloomfield KJ, Hocart CH, Egerton JJG, O’Sullivan OS, Penillard A, Weerasinghe LK, Atkin OK. 2018. Plasticity of photosynthetic heat tolerance in plants adapted to thermally contrasting biomes. Plant, Cell & Environment 41, 1251–1262.

