## Supplementary materials for "Wheat photosystem II heat tolerance responds dynamically to short and long-term warming"

**Table S1.** Pedigree information for wheat genotypes grown for the three field studies and one controlled environment study described in the materials and methods.

| Reference no. | | Pedigree | Note | Group, geographical origin |
| --- | --- | --- | --- | --- |
| **Field studies** | | |  |  |
| 84 | Sokoll/2/Sokoll/ 35888 M 500132 | | Backcross of a hexaploid synthetic derived wheat to a heat tolerant tetraploid *T*. *dicoccum* and a hexaploid type selected | Narrabri, Australia |
| 1132 | PBW550//C80.1/*2Batavia | | Cross of heat tolerant Indian cultivar with rust resistant sources | Pune, India |
| 1683 | PBW343+L24+LR28/Lang | | Same as above | Pune, India |
| 1787 | DBW16/Sunstate | | Same as above | Pune, India |
| 1898 | DBW16/Annuello | | Same as above | Pune, India |
| 1943 | DBW16/Gladius | | Same as above | Pune, India |
| 2062 | ISR 812.8/Carinya (1, sister line) | | Heat tolerant Mexican hexaploid landrace cross to Australian cultivar | Obregon, Mexico |
| 2150 | ISR 812.8/Carinya (2, sister line) | | Same as above | Obregon, Mexico |
| 2219 | Ventura/Ido 637//Ventura | | Low phytate mutant crossed to Australian cultivar - pre-screened for heat tolerance | Narrabri, Australia |
| 2254 | D67.2/P66.270//AE.Squarrosa (320)/3/Cunningham/4/Vorb | | Heat tolerant in Mexico (Ciudad Obregon) and Narrabri, Australia. Origin CGIAR | Obregon, Mexico |
| 2255 | SLVS/Attila//WBLL1*2/3/Gondo/CBRD | | Same as above | Obregon, Mexico |
| 2328 | Sokoll/2/Sokoll/35888 M 500132 | | Backcross of a hexaploid synthetic wheat to a heat tolerant tetraploid *T*. *dicoccum* and a hexaploid type selected | Narrabri, Australia |
| Corack |  | | Commercial Australian cultivar, released in 2012 | Roseworthy, Australia |
| Suntop |  | | Same as above | Roseworthy, Australia |
| Trojan |  | | Commercial Australian cultivar, released in 2013 | Roseworthy, Australia |
| Mace | Wyalkatchem/Stylet//Wyalkatchem | | Commercial Australian cultivar, released in 2008 | Roseworthy, Australia |
| 2475 | Attila/3*BCN//Bav92/3/Tilhi/5/Bav92/3/PRL/Sara//TSI/Vee#5/4/Croc_1/Ae.Squarrosa (224)//2*Opata | | Heat tolerant in Mexico (Ciudad Obregon) and Narrabri. Origin CGIAR | Obregon, Mexico |
| 2355 | Seri 82/Shuha's'//CM85295-0101TOPY-2M-0Y-0M-3Y-0M-0AP | | Same as above | Aleppo, Syria |
| 1964 | DBW14/C80.1/2*SR2 Batavia | | Cross of heat tolerant Indian cultivar with rust resistant sources | Pune, India |
| 1704 | PBW343+L24+LR28/Lang | | Same as above | Pune, India |
| 29 | Berkut/2/Berkut/35883 M500110 | | Backcross of a hexaploid wheat to a heat tolerant tetraploid *T*. *dicoccum* and a hexaploid type selected | Narrabri, Australia |
| 143 | Waxwing*2/Kiritati /3/Waxwing*2/Kiritati /2/ 35888 M 500132 | | Same as above | Narrabri, Australia |
| 2316 | RAC 1192/4/2*Attila/3/Weaver*2/TSC//Weaver | | Heat tolerant hexaploid; good performance in Mexico (Ciudad Obregon) and Narrabri, Australia. Origin CGIAR | Obregon, Mexico |
| **Field studies and controlled environment study** | | | |  |
| 2267 | Hubara-8///Mon's'/Ald's'//Bow's' | | Heat tolerant hexaploid; good performance in Sudan and Narrabri. Origin CGIAR | Gezira, Sudan |

**Table S2.** Analysis of variance of factors influencing wheat *T*_crit_ at two Australian field sites

|  | **Genotype** | | **Time of day** | | **Genotype x Time of day** | |
| --- | --- | --- | --- | --- | --- | --- |
|  | d.f. | *F* value | d.f. | *F* value | d.f. | *F* value |
| Dingwall, Victoria | 5 | 3.7 ^**^ | 3 | 15.8 ^***^ | 15 | 2 ^*^ |
|  | **Genotype** | | **Phenological stage** | | **Genotype x Phenological stage** | |
|  | d.f. | *F* value | d.f. | *F* value | d.f. | *F* value |
| Barraport West, Victoria | 3 | 5.0** | 2 | 14.9 ^***^ | 6 | 2.2 ^*^ |

^*^*P* < 0.05; ^**^*P* < 0.01; ^***^*P* < 0.001. Six out of the 20 genotypes sown at Dingwall in 2017 were sampled every six hours to measure diel variation in *T*_crit_ over the course of a day. Four of the 20 genotypes sown at Barraport West in 2018 were sampled from all three time of sowing plots to measure variation in wheat *T*_crit_ at varying phenological stages.

**Table S3.** Analysis of variance of effect of time of sowing and genotype on wheat *T*_crit_ at three Australian field sites

|  | **Time of sowing** | | **Genotype** | | **Time of sowing x Genotype** | |
| --- | --- | --- | --- | --- | --- | --- |
|  | d.f. | *F* value | d.f. | *F* value | d.f. | *F* value |
| Dingwall, Victoria | 2 | 23.1 ^***^ | 19 | 0.5 ^ns^ | 38 | 0.9 ^ns^ |
| Barraport West, Victoria | 2 | 62.1 ^***^ | 19 | 2.1 ^**^ | 38 | 2 ^**^ |
| Narrabri, New South Wales | 1 | 13.7 ^***^ | 23 | 2.3 ^***^ | 23 | 1 ^ns^ |

^*^*P* < 0.05; ^**^*P* < 0.01; ^***^*P* < 0.001; ^ns^ = not significant. The same 20 genotypes that were sown in Dingwall and Barraport West were also sown at Narrabri in 2019, along with an additional four genotypes.

**Table S4.** Analysis of variance of effect of time of sowing and genotype origin on wheat *T*_crit_ at three Australian field sites

|  | **Time of sowing** | | **Genotype origin** | | **Time of sowing x Genotype origin** | |
| --- | --- | --- | --- | --- | --- | --- |
|  | d.f. | *F* value | d.f. | *F* value | d.f. | *F* value |
| Dingwall, Victoria | 2 | 15.2 ^***^ | 4 | 0.2 ^ns^ | 8 | 0.7 ^ns^ |
| Barraport West, Victoria | 2 | 42.2 ^***^ | 4 | 2.5 ^*^ | 8 | 1 ^ns^ |
| Narrabri, New South Wales | 1 | 4.5 ^*^ | 5 | 2.5 ^*^ | 5 | 0.7 ^ns^ |

^*^*P* < 0.05; ^**^*P* < 0.01; ^***^*P* < 0.001; ^ns^ = not significant. The same 20 genotypes that were sown in Dingwall and Barraport West were also sown at Narrabri in 2019, along with an additional four genotypes.
